# Core Genes Evolve Rapidly in the Long-term Evolution Experiment with *Escherichia coli*

**DOI:** 10.1101/042069

**Authors:** Rohan Maddamsetti, Philip J. Hatcher, Anna G. Green, Barry L. Williams, Debora S. Marks, Richard E. Lenski

## Abstract

Bacteria can evolve rapidly under positive selection owing to their vast numbers, allowing their genes to diversify by adapting to different environments. We asked whether the same genes that are fast evolving in the long-term evolution experiment with *Escherichia coli* (LTEE) have also diversified extensively in nature. We identified ~2000 core genes shared among 60 *E. coli* strains. During the LTEE, core genes accumulated significantly more nonsynonymous mutations than flexible (i.e., noncore) genes. Furthermore, core genes under positive selection in the LTEE are more conserved in nature than the average core gene. In some cases, adaptive mutations appear to fine-tune protein functions, rather than merely knocking them out. The LTEE conditions are novel for *E. coli*, at least in relation to the long sweep of its evolution in nature. The constancy and simplicity of the environment likely favor the complete loss of some unused functions and the fine-tuning of others.

**Competing Interests Statement:** We, the authors, declare that we have no conflicts of interest.

## Introduction

By combining experimental evolution and genomic technologies, researchers can study in fine detail the genetic underpinnings of adaptation in the laboratory (Barrick and Lenski 2013). However, questions remain about how the genetic basis of adaptation might differ between experimental and natural populations (Bailey and Bataillon 2016).

To address that issue, we examined whether the genes that evolve most rapidly in the long-term evolution experiment (LTEE) with *Escherichia coli* also evolve and diversify faster than typical genes in nature. If so, the genes involved in adaptation in the LTEE might also be involved in local adaptation to diverse environments in nature. On the other hand, it might be the case that the genes involved in adaptation during the LTEE diversify more slowly in nature than typical genes. Perhaps these genes are highly constrained in nature by purifying selection. For example, they may play important roles in balancing competing metabolic demands or fluctuating selective pressures in the complex and variable natural world, but they can be optimized to fit the simplified and stable conditions of the LTEE.

To test these alternative hypotheses, we compare the signal of positive selection across genes in the LTEE to the sequence diversity in a set of 60 clinical, environmental, and laboratory strains of *E. coli*—henceforth, the “*E. coli* collection”—and to the divergence between *E. coli* and *Salmonella enterica* genomes, respectively. We find that the genes that have evolved the fastest in the LTEE, based on parallel nonsynonymous mutations that are indicative of positive selection, tend to be conserved core genes in the *E. coli* collection. We can exclude recurrent selective sweeps at these loci in nature as an explanation for their limited diversity because the genes and the particular amino-acid residues under positive selection in the LTEE have diverged slowly since the *Escherichia*–*Salmonella* split. We also present structural evidence that some of the nonsynonymous mutations—especially those where identical amino-acid changes evolved in parallel—are beneficial because they fine-tune protein functions, rather than knocking them out.

## Results

### Core Genes Are Functionally Important

To make consistent comparisons between the LTEE lines and the *E. coli* collection, we analyzed single-copy genes with homologs in all 60 sequenced genomes in the *E. coli* collection. For the purpose of our study, we define this set of panorthologous genes as the *E. coli* core genome and the set of all other genes as the flexible genome (Materials and Methods). We used published data from the Keio collection of single-gene knockouts in *E. coli* K-12 (Baba *et al.* 2006) to test whether the core genes tend to be functionally more important than the flexible genes based on essentiality and growth yield. As expected, core genes are indeed more essential than flexible genes (Welch’s *t* = 6.60, d.f. = 3387.8, one-tailed *p* < 10^−10^), and knockouts of core genes cause larger growth-yield defects than do knockouts of flexible genes in both rich (Welch’s *t* = 3.79, d.f. = 3379, one-tailed *p* < 0.0001) and minimal media (Welch’s *t* = 4.95, d.f. = 3457.3, one-tailed *p* < 10^−6^).

### Core Genes Evolve Faster than Flexible Genes in the LTEE

We first examined the mutations in single genomes sampled from each of the 12 LTEE populations after 50,000 generations. Six of these populations evolved greatly elevated point-mutation rates at various times during the LTEE (Sniegowski *et al.* 1997, Blount *et al.* 2012, Wielgoss *et al.* 2013, Tenaillon *et al.* 2016). As a consequence of their much higher mutation rates, a much larger fraction of the mutations seen in hypermutable populations are expected to be neutral or even deleterious passengers (hitchhikers), as opposed to beneficial drivers, in comparison to those populations that retained the low ancestral point-mutation rate (Tenaillon *et al.* 2016). In genomes from the nonmutator populations, we observe a highly significant excess of nonsynonymous mutations in the core genes. Specifically, the core genes constitute ~48.5% of the total coding sequence in the genome of the ancestral strain, but ~71% (123/174) of the nonsynonymous mutations in the 50,000-generation clones are found in the core genes (Table 1, row 1, *p* < 10^−8^). However, there is no significant difference in essentiality between the core genes with zero versus one or more nonsynonymous mutations (Welch’s *t* = 1.56, d.f. = 180.6, two-tailed *p* = 0.1204), so there is no evidence that core genes evolving in the LTEE are either enriched or depleted for essential genes.

**Table 1.**
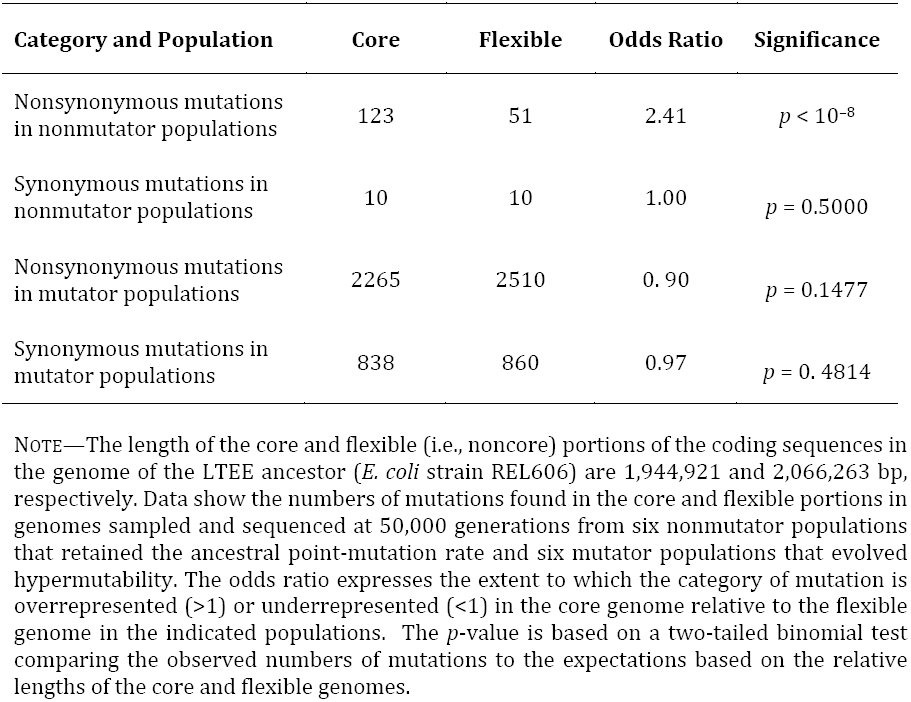
Nonsynonymous mutations are overrepresented in the core genome of nonmutator LTEE populations.

By contrast, the frequency of synonymous mutations does not differ significantly between the core and flexible genes (Table 1, row 2), demonstrating that the excess of nonsynonymous mutations in core genes is driven by selection, not by their propensity to mutate (Maddamsetti *et al*. 2015). Also, the frequencies of both nonsynonymous and synonymous mutations in core versus flexible genes are close to the null expectations in the populations that evolved hypermutability (Table 1, rows 3 and 4).

These results show that core genes are evolving faster, on average, than the flexible noncore genome in the LTEE populations that retained the ancestral point-mutation rate. This faster evolution is consistent with some subset of the core genes being under positive selection to change from their ancestral state during the LTEE. We then wanted to know how the rates of evolution of core genes observed in the LTEE compare to the rates of evolution of the same genes over the longer timescale of *E. coli* evolution. We used the *G* scores from Tenaillon *et al.* (2016) as a measure of the rate of evolution of each core gene in the LTEE. The *G*-score statistic expresses the excess number of independent nonsynonymous mutations in the nonhypermutable lineages relative to the number expected, given the length of that gene’s coding sequence (relative to all coding sequences) and the total number of such mutations. To measure the rate of evolution of each core gene in nature, we used Nei’s diversity metric (Nei and Li 1979); in brief, we calculate the average number of differences per site among all pairs of sequences in the core gene alignments.

We found a very weak, albeit significant, negative correlation between a core gene’s *G* score in the LTEE and its diversity in the *E. coli* collection (Spearman-rank correlation *r* = −0.0701, two-tailed *p* = 0.0019; Fig. 1A). Only 163 genes in the core genome had positive *G* scores (i.e., one or more nonsynonymous mutations in nonhypermutable lineages) in the LTEE, and we do not find a significant correlation between the *G* score and sequence diversity using only those genes (Spearman-rank correlation *r* = −0.0476, two-tailed *p* = 0.5463; Fig. 1B). However, taken together, the 163 core genes with positive *G* scores have significantly lower diversity in the *E. coli* collection than do the 1805 with zero *G* scores (Mann-Whitney U = 125,660, two-tailed *p* = 0.0020; Fig. 1C). Hence, the difference between the core genes with and without nonsynonymous mutations in the nonmutator lineages drives the weak overall negative correlation.

**FIG. 1.**
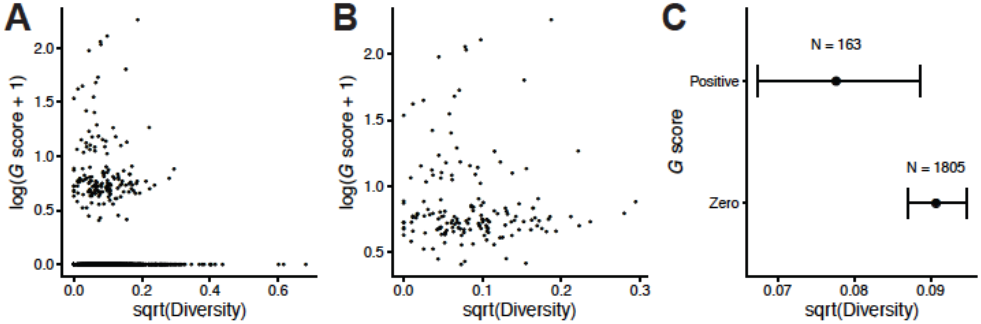
Relationship between positive selection in the LTEE and nonsynonymous sequence diversity of core genes in the *E. coli* collection of 60 clinical, environmental, and laboratory strains. The *G* score provides a measure of positive selection based on the excess of nonsynonymous mutations in the LTEE lineages that retained the ancestral point-mutation rate. The log_10_ and square-root transformations of the *G* score and sequence diversity, respectively, improve visual dispersion of the data for individual genes, but they do not affect the nonparametric tests performed, which depend only on rank order. (A) *G* scores and sequence diversity are very weakly negatively correlated across all 1968 core genes (Spearman-rank correlation *r* = −0.0701, *p* = 0.0019). (B) The correlation is not significant using only the 163 genes with positive *G* scores (Spearman-rank correlation *r* = −0.0476, *p* 11 = 0.5463). (C) The 163 core genes with positive *G* scores in the LTEE have significantly lower nonsynonymous sequence diversity in natural isolates than the 1805 genes with zero *G* scores (Mann-Whitney *U* = 125,660, *p* = 0.0020). Error bars show 95% confidence intervals around the median.

By using segregating polymorphisms in the *E. coli* collection, our measure of the rate of evolution of core genes in nature might be dominated by transient variation or local adaptation. By contrast, core genes found in different species have diverged over a longer timescale and should be less affected by these issues. Therefore, we repeated the above analyses using the set of 2853 panorthologs—single-copy genes that map one-to-one across species (Lerat *et al.* 2003, Cooper *et al.* 2010)—for *E. coli* and *Salmonella enterica*. We found a weak but again significant negative correlation across genes between their *G* scores in the LTEE and interspecific divergence (Spearman rank-correlation *r* = −0.0911, two-tailed *p* < 10^−5^; Fig. 2A). This negative correlation remains significant even if we consider only those 210 panorthologs with positive *G* scores in the LTEE (Spearman rank-correlation *r* = −0.2564, two-tailed *p* = 0.0002; Fig. 2B). In addition, the panorthologs with positive *G* scores are less diverged between *E. coli* and *S. enterica* than the 2643 panorthologs with zero *G* scores (Mann-Whitney U = 223,330, two-tailed *p* < 10^−5^; Fig. 2C).

**FIG. 2.**
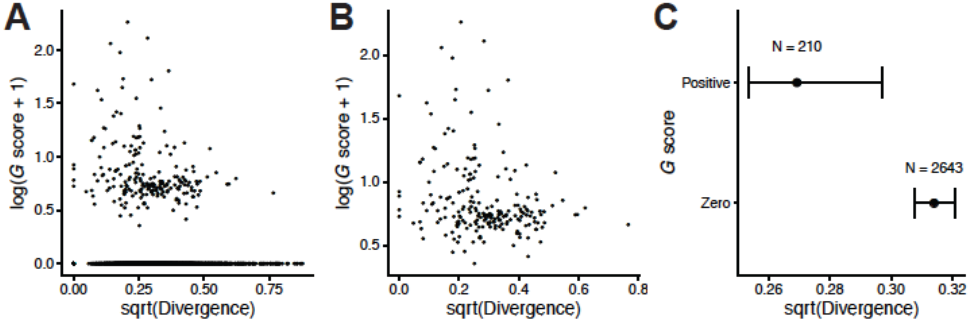
Relationship between positive selection in the LTEE and nonsynonymous sequence divergence of panorthologs between *E. coli* (strain REL606) and *S. enterica*. REL606 is the common ancestor of the LTEE populations. See Fig. 1 for additional details. (A) *G* scores and divergence are negatively correlated across all 2853 panorthologs (Spearman-rank correlation *r* = −0.0911, *p* < 10^−5^). (B) The correlation remains significant even when using only the 210 panorthologs with positive *G* scores (Spearman-rank correlation *r* = −0.2564, *p* = 0.0002). (C) The 210 panorthologs with positive *G* scores in the LTEE are significantly less diverged between *E. coli* and *S. enterica* in natural isolates than the 2643 panorthologs with zero *G* scores (Mann-Whitney *U* = 223,330, *p* = < 10^−5^). Error bars show 95% confidence intervals around the median.

In sum, these analyses contradict the hypothesis that those genes that have evolved fastest in the LTEE are ones that also evolve and diversify faster than typical genes in nature. Instead, they support the hypothesis that the genes involved in adaptation during the LTEE tend to be more conserved than typical genes in nature, presumably because they are constrained in nature by purifying selection. When the bacteria evolve under the simple and stable ecological conditions of the LTEE, these previously conserved genes evolve in and adapt to their new environment.

### Protein Residues that Changed in the LTEE Are Also Conserved in Nature

It is possible that the mutations in the LTEE occurred at highly variable sites in otherwise conserved proteins. To examine this issue, we asked whether the nonsynonymous changes found in the nonmutator LTEE lineages at 50,000 generations tended to occur in fast-evolving codons. For the 96 proteins with such mutations in the LTEE, we calculated the diversity at the mutated sites and in the rest of the protein for the 60 genomes in the *E. coli* collection. The sites that had changed in the LTEE were significantly less variable than the rest of the protein in that collection (Wilcoxon signed-rank test, *p* < 10^−7^). In fact, only 7 of these 96 proteins had any variability at those sites in the *E. coli* collection, and these 7 proteins account for only 9 of the 141 amino-acid mutations in the 96 proteins. We obtained similar results based on the divergence between *E. coli* and *Salmonella.* In the 50,000-generation LTEE clones, 144 nonsynonymous mutations occurred in 102 panorthologous genes, and only 6 of the mutations were at diverged sites. These results show that the particular residues under positive selection in the LTEE are, in fact, ones that tend to be conserved in nature, even over the ~100 million years since *Escherichia* and *Salmonella* diverged (Ochman *et al.* 1999).

### Knockout Versus Fine-tuning Beneficial Mutations in the LTEE

How did the mutations that fixed in the LTEE drive adaptation to the bacteria’s new environment? In some cases, these beneficial mutations might fine-tune protein function, whereas in other cases (e.g., deletions) they might be beneficial by knocking out the protein function. We examined two lines of evidence for fine-tuning mutations: gene essentiality, because essential genes cannot be knocked out; and parallel evolution at the amino-acid level, because we do not expect strong molecular constraints if many different mutations can knock out a gene’s function. To address the second issue, we necessarily restricted the analysis to the 57 genes with two or more nonsynonymous changes in nonmutator lineages. We used the same 57 genes to address the first issue for consistency, and because genes with multiple nonsynonymous changes are evidently under positive selection in the LTEE.

#### Evidence for fine-tuning based on essentiality

We would expect essential genes that were under positive selection in the LTEE to have mutations that fine-tune protein function, not knockout mutations that eliminate an essential function. KEIO essentiality scores range from +3 to –4, where more positive scores indicate essentiality and more negative scores indicate dispensability (Fig. 3). We labeled the 57 genes under strong positive selection according to the presence of possible knockout mutations in nonmutator genomes—that is, small indels that often disrupt the reading frame, IS-element insertions, and large deletions affecting the gene. The 16 genes with positive essentiality scores are less likely to have been impacted by possible knockout mutations than the 40 genes with negative scores (Fisher’s exact test: one-tailed *p* = 0.0297); one gene has a score of zero. Of the genes with positive essentiality scores, only *topA* and *mrdA* had any possible knockout mutations in any of the sequenced nonmutator genomes. The candidate knockout in *topA* is a small indel found in only one genome (one of two clones from population Ara-6 at generation 50,000). This mutation causes a frameshift in the penultimate codon of the gene, adding 3 amino acids to the tail of the protein before reaching a new stop codon. Therefore, this small indel in *topA* probably does not destroy the gene’s function. In the second case, there is a large (161,226 bp) deletion in all Ara+1 clones from 30,000 generations onward that removed *mrdA* and many other genes. This *mrdA* mutation is clearly a true knockout. Also, the *G*-scores for gene-level parallelism included nonsense mutations with nonsynonymous mutations, whereas some nonsense mutations might also be knockouts. However, only 5 of the 57 genes, all with negative essentiality scores (*yijC*, *malT, yabB, ybaL,* and *yeeF*), have nonsense mutations in any of the nonmutator genomes, and 4 of them (all except *yijC*) are also affected by small indels or large deletions. This line of evidence therefore supports the hypothesis that some of the beneficial mutations in the LTEE modulate protein function, rather than knocking it out.

**FIG. 3.**
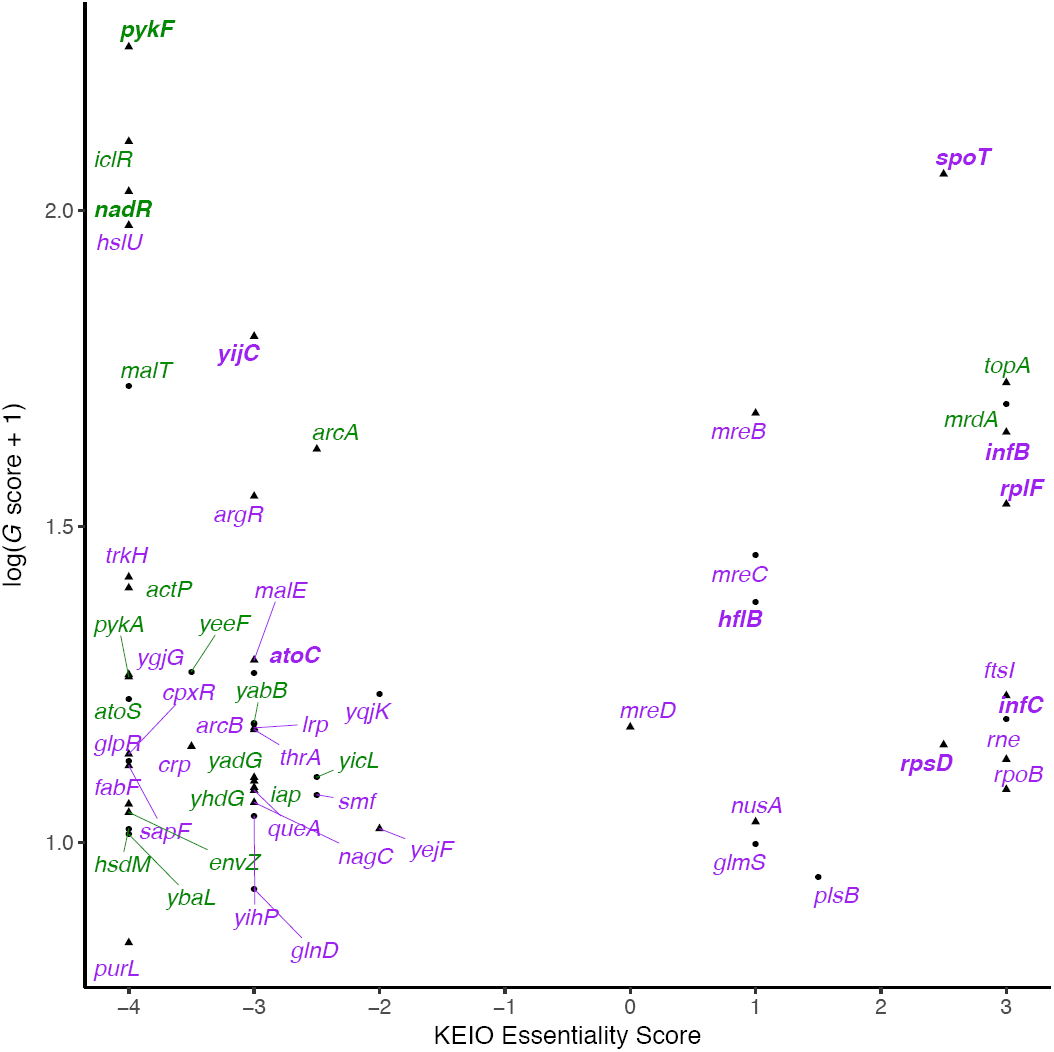
KEIO essentiality score and *G* score for the 57 genes with 2 or more nonsynonymous changes in nonmutator LTEE genomes. The transformation of the *G* score improves visual dispersion of the data for clarity. Triangles are core genes (panorthologs) and circles are noncore flexible genes. Genes affected by at least one potential knockout mutation (small indel, IS-element insertion, or large deletion) are labeled in green, and genes without any of these potential knockout mutations in purple. Also, 10 genes that had parallel mutations at the amino-acid level are additionally indicated in bold.

#### Evidence for fine-tuning based on parallelism at the amino-acid level

The second line of evidence for fine-tuning (rather than complete loss of function) involves genes in which the same amino-acid mutations evolved in multiple populations. If mutations in a particular gene were beneficial because they knocked out the protein function, then we would expect to find many different mutations in those genes, including varied nonsynonymous changes (as well as indels). By contrast, parallel evolution at the amino-acid level would imply that those specific changes to the protein were more beneficial than other possible mutations.

Ten of the 57 genes that showed gene-level parallel evolution also show parallelism at the amino-acid level. We mapped these mutations onto the three-dimensional structures of the protein in *E. coli* or close homologs (Berman *et al.* 2000). In 8 of these 10 cases, the parallel mutations in the proteins are within 8 ångströms of a bound protein or ligand (Fig. 4). In the other 2 cases, the entire protein is in close contact with ribosomal RNA.

**FIG. 4.**
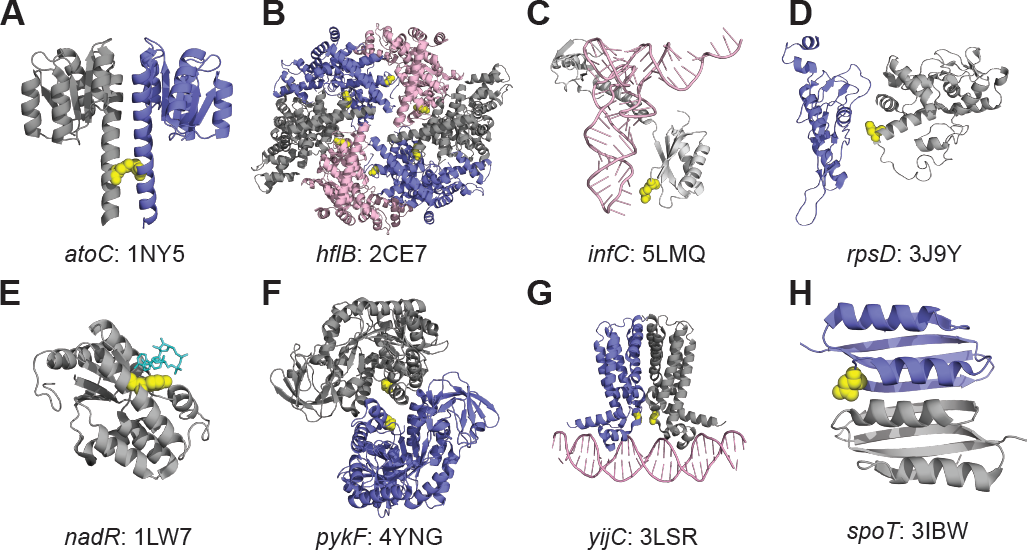
Parallel amino-acid mutations in the LTEE occur at protein interfaces. For clarity, only relevant protein domains are shown. (A) The I129S mutation is on the dimerization interface of the response regulator AtoC, based on the *Aquifex aeolicus* structure 1NY5. (B) Q506L occurs on a multimerization interface of the metalloprotease FtsH, encoded by *hflB*, in the *Thermotoga maritima* structure 2CE7. (C) R132C in ribosomal initiation factor IF3 interacts with the anticodon of the fMet-tRNA in the *E. coli* structure 5LMQ. (D) Mutations at residue 50 in 30S ribosomal protein S4 lie on the interface with protein S5 in the *E. coli* ribosome structure 3J9Y. (E) Mutations at residue 294 directly contact the coenzyme NAD in the *Haemophilus influenzae* NadR protein structure 1LW7, while mutations at residues 290 and 298 are adjacent to 294 on the same face of the alpha helix. (F) A301S occurs at the A/A’ multimerization interface of pyruvate kinase, encoded by *pykF*, in the *E. coli* structure 4YNG. (G) T30N occurs at the dimerization interface of the DNA-binding domain of the transcriptional repressor FabR in the *Pseudomonas aeruginosa* structure 3LSR. (H) N653H occurs at the dimerization interface between ACT4 amino-acid binding domains of the bifunctional (p)ppGpp synthase/hydrolase SpoT in the *Chlorobium tepidum* structure 3IBW.

In *atoC*, an I129S mutation occurs in both 50,000-generation clones from population Ara+1 and in a 30,000-generation clone from Ara+4; this mutation maps to a dimerization interface (Fig. 4A). We found a Q506L mutation in *hflB* in both 50,000-generation clones from Ara+5 and in a 30,000-generation clone from Ara–5; this mutation maps to a multimerization interface (Fig. 4B). In *infC,* an R132C substitution was fixed on the line of descent in both the Ara+4 and Ara-6 populations (Fig. 4C). This mutation interacts with the anticodon of the fMet-tRNA during translation initiation on the ribosome. A D50G mutation in *rpsD* is on the line of descent (i.e., reached fixation or nearly so) in five populations: Ara– 1, Ara–2, Ara–4, Ara+2, and Ara+3. This particular residue of the 30S ribosomal protein S4 has the strongest signal among all S4 residues for interaction with the ribosomal protein S5 in a maximum-entropy model of sequence variation within and between the two proteins
 (Hopf *et al*. 2014), and the mutation clearly maps to their interface (Fig. 4D). The R132C *infC* mutation and the D50G *rpsD* mutation are the only nonsynonymous mutations found in these genes in any of the sequenced LTEE genomes (including even those that evolved hypermutability), providing further evidence that their benefits result from specific fine-tuning effects.

In *nadR*, *pykF*, and *yijC* (*fabR*), we also found parallel evolution at the amino-acid level across some LTEE populations, but with knockout mutations in other populations. In the case of *nadR*, Y294C substitutions are on the line of descent in four populations: Ara–5, Ara+2, Ara+4, and Ara+5. This mutation interacts with NAD in a homologous protein structure; mutations at nearby residues 290 and 298 occurred in two other populations, and these residues are on the same face of the alpha helix (because alpha helices have a period of 3.6 residues) that interacts with NAD (Fig. 4E). In fact, nonsynonymous mutations in *nadR* fixed in all but one of the 12 LTEE populations; the exception was population Ara+1, in which an IS*150* element inserted into the gene. In the case of *pykF*, an A301S mutation fixed on the line of descent in three populations: Ara–5, Ara+1, and Ara+5. This residue lies at the A/A’ multimerization interface of the *pykF* tetramer (Fig. 4F), which is implicated in allostery in response to fructose 1,6-biphosphate binding (Donovan *et al.* 2016). However, *pykF* knockout mutations are also beneficial in the LTEE environment (Barrick *et al.* 2009, Khan *et al.* 2011). A 1-bp deletion fixed in Ara+4, and many *pykF* mutations that cause frameshifts are found off the line of descent in many LTEE populations, including disruption by IS*150* transposons. Biochemical analyses indicate that *pykF* alleles vary in their catalytic and allosteric properties, so that some changes in function might be more beneficial than knockouts of the protein (R. Dobson and T. Cooper, personal communication, January 2017). We also find parallelism at the amino-acid level in *yijC* (*fabR*). A T30N mutation is found off the line of descent in a 500-generation clone from Ara–1 clone, a 500-generation clone from Ara–2, and a 1500-generation clone from Ara+1. Other early mutations at this locus were found in populations Ara+1, Ara+5, Ara–3, and Ara–6, including a Q172* nonsense mutation that persisted in Ara–3 for at least 1000 generations. In no case, however, did any mutation in *yijC* (*fabR)* become fixed in the LTEE, indicating that positive selection was insufficient to drive them to fixation. Structural analysis shows the T30N mutation is at the dimerization interface of the protein on its DNA-binding domain (Fig. 4G). We also examined the F83V mutation in the 50S ribosomal protein L6, which is encoded by *rplF*, as well as the K717E mutation in translation initiation factor IF-2, encoded by *infB.* In both of these cases, the entire protein is in close contact with ribosomal RNA.

In contrast to these cases of parallel evolution at the amino-acid level, different nonsynonymous mutations in the *spoT* gene were fixed in populations Ara–1, Ara–2, Ara–4, Ara–6, Ara+2, Ara+4, and Ara+6. These mutations affect different domains of the SpoT protein (Ostrowski *et al*. 2008). The complete absence of any putative knockout mutations at this locus across the LTEE (Tenaillon *et al.* 2016) indicates that mutations in *spoT* are probably functional, and not knockouts. Further evidence of fine-tuning evolution in *spoT* is the fact that an N653H mutation evolved twice, being present in an Ara+3 2000-generation clone and an Ara–5 30,000-generation clone, although in neither case did it fix. Structural analysis shows that this mutation lies at the dimerization interface of an ACT4 domain (Fig. 4H). In general, ACT domains are involved in allosteric control in response to amino-acid binding (Cross *et al.* 2013).

## Discussion

It has been long known that, in nature, some genes evolve faster than others. In most cases, the more slowly evolving genes are core genes—ones possessed by most or all members of a species or higher taxon—and their sequence conservation reflects constraints that limit the potential for the encoded proteins to change while retaining their functionality. As a consequence, the ratio of nonsynonymous to synonymous mutations also tends to be low in these core genes. By contrast, we found that nonsynonymous mutations in nonmutator lineages of the LTEE occurred disproportionately in the core genes shared by all *E. coli* (Table 1). Moreover, even among the core genes, those that experienced positive selection to change in the LTEE are both less diverse over the *E. coli* species (Fig. 1) and less diverged between *E. coli* and *S. enterica* (Fig. 2) than other core genes. Also, the specific sites where mutations arose in the LTEE are usually more conserved than the rest of the corresponding protein, thus excluding the possibility that mutations occurred at a subset of fast-evolving sites in otherwise slow-evolving genes.

In fact, many of the core genes under selection in the LTEE perform vital functions or regulate key aspects of cell physiology (Table 2). In comparison to their *E. coli* B ancestor, the evolved bacteria have a shorter lag phase when transferred into fresh medium, a higher maximum growth rate, improved glucose transport, larger cell size, and altered cell shape (Lenski *et al.* 1998). Glucose transport is probably improved, in part, by mutations in genes encoding the pyruvate kinases that catalyze the phosphorylation of phosphoenolpyruvate (PEP) to pyruvate. By inhibiting the forward reaction, or perhaps promoting the reverse reaction, mutations affecting the kinases would increase the concentration of PEP, which drives the phosphotransferase system that brings glucose into the cell (Woods *et al.* 2006). Global regulatory networks also have evolved in the LTEE (Cooper *et al.* 2003; Philippe *et al.* 2007). DNA superhelicity, which links chromosome structure to gene regulation, has been under strong selection in the LTEE (Crozat *et al.* 2005; Crozat *et al.* 2010), as has the CRP regulon that coordinates metabolism with cellular protein production (You *et al.* 2013) and the ppGpp regulon that regulates ribosome synthesis in response to levels of available amino acids in the cell (Scott *et al.* 2010; Scott *et al.* 2014).

**Table 2.**
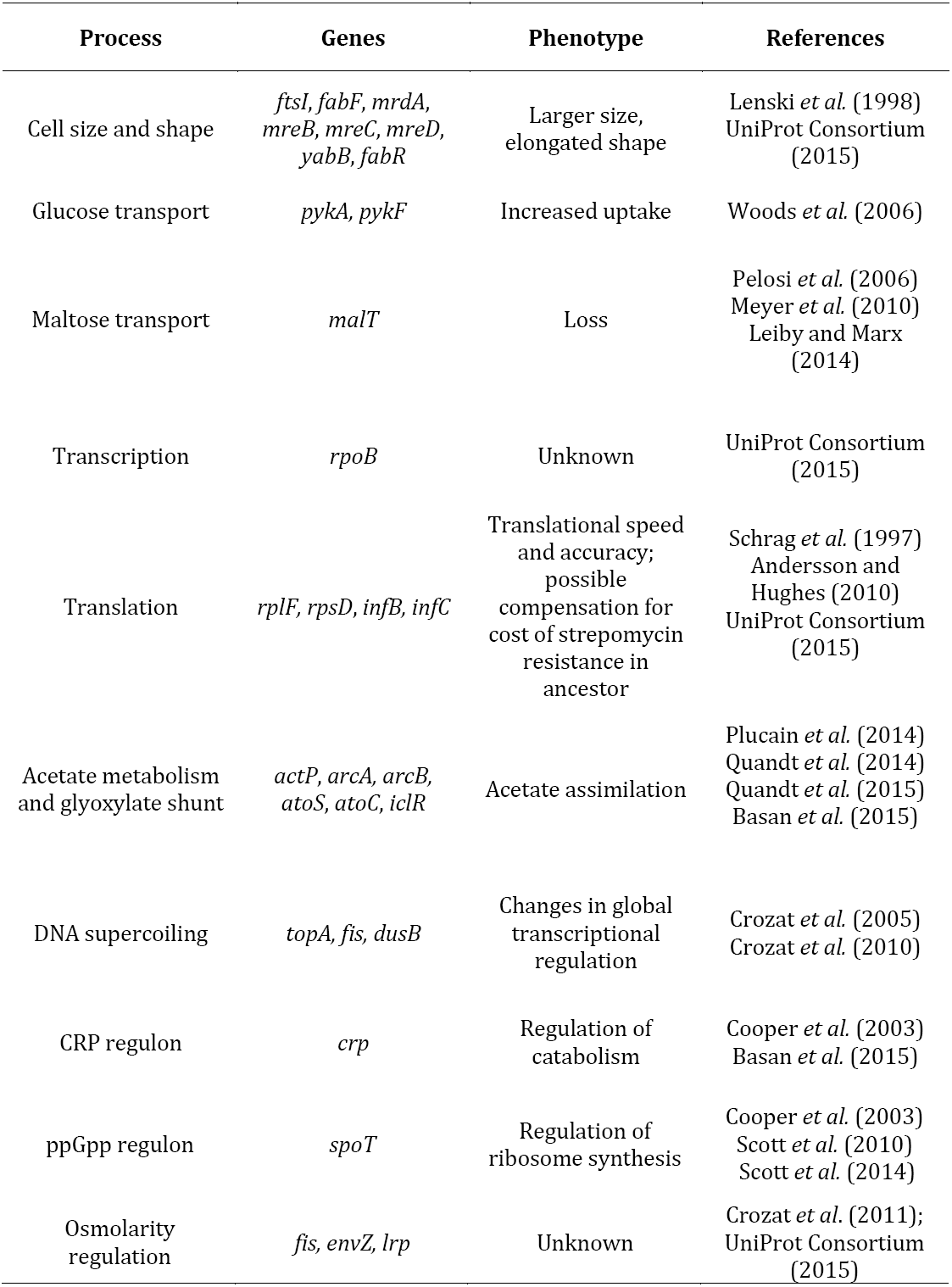
Genes and their associated phenotypes that show evidence of positive selection in the LTEE. See also Tenaillon *et al.* (2016) for evidence of gene-level parallelism.

Some other mutations may ameliorate the fitness cost associated with a mutation in *rpsL* that confers resistance to strepomycin in the ancestral strain REL606, which was selected prior to the LTEE (Studier *et al.* 2009). The LTEE populations have maintained this resistance despite 50,000 generations of relaxed selection, probably owing to the fixation of compensatory mutations in ribosomal protein-encoding genes such as *rpsD* (Schrag *et al.* 1997, Andersson and Hughes 2010). Researchers have long known that mutations at the interface of ribosomal proteins S4 and S5 can compensate for streptomycin resistance, and that such mutations can affect translational speed and accuracy (Agarwal *et al.* 2015). That context, in addition to our new finding that the R132C mutation in *infC* interacts with fMet-tRNA during translation initiation, provides evidence for positive selection on translational speed, accuracy, or both in the LTEE.

It is clear, then, that the specific conditions of the LTEE have favored new alleles in core genes that are usually highly conserved in nature. From one perspective, this result is surprising: the LTEE’s 37°C temperature is typical for humans and many other mammalian bodies where *E. coli* lives; the limiting resource is glucose, which is *E. coli*’s preferred energy source, such that it will repress the expression of genes used to catabolize other resources when glucose is available; and the LTEE does not impose other stressors such as pH, antibiotics, or the like. However, the very simplicity and constancy of the LTEE are presumably novel, or at least atypical, in the long sweep of *E. coli* evolution (Fig. 5). In other words, that uniformity and simplicity—including the absence of competitors and parasites as well as host-dependent factors—stand in stark contrast to the variable and complex communities that are *E. coli*’s natural habitat (Blount 2015). Of course, evolutionary outcomes depend on the environment and the constraints it imposes. For example, compensatory mutations in *rpsD* and *rpsE* readily evolved when streptomycin-resistant *Salmonella* populations were passaged in broth, but not when they were passaged in mice (Björkman *et al.* 2000).

**FIG. 5.**
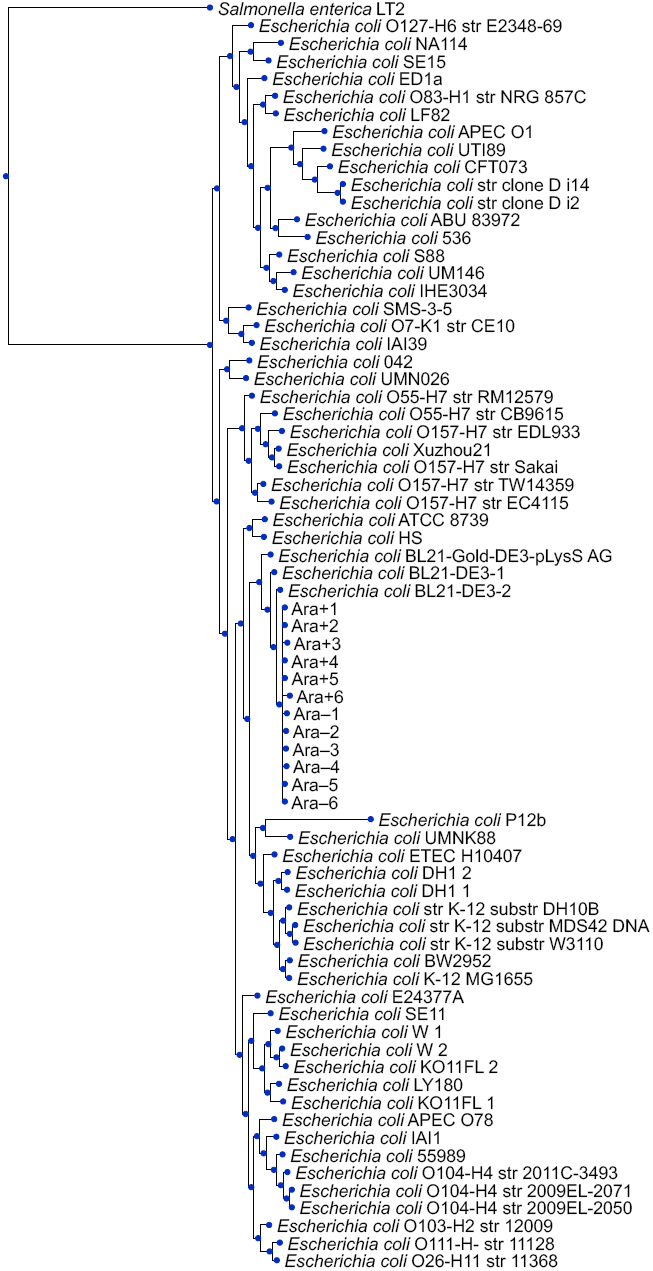
Neighbor-joining evolutionary tree based on the genomes analyzed in this study. They include 60 clinical, environmental, and laboratory strains in the *E. coli* collection, 12 50,000-generation clones from the LTEE (labeled Ara+1 to Ara–6), and *Salmonella enterica*.

Given the importance and even essentiality of many core genes, it seems unlikely that the beneficial nonsynonymous mutations in the LTEE only cause complete losses of function. Indeed, many of these beneficial mutations appear to fine-tune the regulation and expression of functions that contribute to the bacteria’s competitiveness and growth in the simple and predictable environment of the LTEE (Table 2, Fig. 4). As further evidence, the functional effects and fitness consequences of some of the evolved alleles depend on earlier mutations in other genes. Thus, different alleles at the same locus that evolved in different lineages may have different effects, and particular combinations of alleles are sometimes necessary to confer a selective advantage. Such epistasis between evolved alleles has been demonstrated in several LTEE populations and involves various genes including *spoT* and *topA* in population Ara–1 (Woods *et al.* 2011); *arcA, gntR,* and *spoT* in Ara–2 (Plucain *et al.* 2014); and *citT*, *dctA*, and *gltA* in Ara–3 (Quandt *et al.* 2014, Quandt *et al.* 2015). In the case of Ara–3, these epistatic interactions were important for that population’s novel ability to grow on citrate in the presence of oxygen (Blount *et al.* 2008, Blount *et al.* 2012).

By contrast, some other genes that were repeatedly mutated in the LTEE—not by point mutations, but instead by deletions and transposable-element insertions—typically encode noncore, nonessential functions including prophage remnants, plasmid-derived toxin-antitoxin modules, and production of surface structures that are probably important for host colonization (Tenaillon *et al.* 2016). Both types of change have been shown to be adaptive in the LTEE environment—point mutations by affecting a gene’s function and the expression of interacting genes (Cooper *et al.* 2003, Philippe *et al.* 2007), and indels by eliminating unused and potentially costly functions (Cooper *et al.* 2001).

Of course, other evolution experiments may well produce different types of genomic changes, including in some cases perhaps a preponderance of point mutations in noncore genes. For example, if the experimental environment involves lethal agents such as phages or antibiotics, then perhaps only a few noncore genes might be the targets of selection, and the resulting mutations might even interfere with adaptation to other aspects of the environment (Scanlan *et al.* 2015). Similarly, adaptation to use novel resources—such as the ability to use the citrate that has been present throughout the LTEE, but which only one population has discovered how to exploit (Blount *et al.* 2008, Blount et al. 2012)—may produce a different genetic signature. Yet other signatures might emerge if horizontal gene transfer from other strains or species provided a source of variation (Souza *et al.* 1997). Imagine, for example, a scenario in which gene flow allowed *E. coli* to obtain DNA from a diverse natural community; in that case, a transporter acquired from another species might well provide an easier pathway to use the citrate in the LTEE environment.

We can turn the question around from asking why core genes evolve so quickly in the LTEE, to why they usually evolve slowly in nature. Core genes encode functions that, by definition, are widely shared, and so their sequences have had substantial time to diverge and become fine-tuned to different niches (Biller *et al.* 2015). As a consequence, there are fewer opportunities for new alleles of core genes to provide an advantage. Moreover, given the diversity of species (including transients) in most natural communities, extant species usually fill any vacant niches that might appear as a result of environmental changes faster than *de novo* evolution. Nonetheless, mutations in conserved core genes might sometimes provide the best available paths for adaptation to new conditions, such as when formerly free-living or commensal bacteria become pathogens (Lieberman *et al.* 2011). In such cases, finding parallel or convergent changes offers a way to identify adaptive mutations when they occur in core genes. For example, *E. coli* and *S. enterica* have been found to undergo convergent changes at the amino-acid level in core genes when strains evolve pathogenic lifestyles (Chattopadhyay *et al.* 2009; Chattopadhyay *et al.* 2012).

In summary, the genetic signatures of adaptation vary depending on circumstances including the novelty of the environment from the perspective of the evolving population, the complexity of the biological community in which the population exists, the intensity of selection, and the number and types of genes that can produce useful phenotypes. In the LTEE, nonsynonymous mutations in core genes that encode conserved and even essential functions for *E. coli* have provided an important source of the fitness gains in the evolving populations over many thousands of generations (Wiser *et al.* 2013, Lenski *et al.* 2015).

## Materials and Methods

### Panortholog Identification in the *E. coli* Collection

We downloaded the nucleotide and amino-acid sequences from GenBank for 60 fully sequenced *E. coli* genome accessions (Table S1). We refer to this diverse set of clinical, environmental, and laboratory strains as the *E. coli* collection. We identified 1968 single-copy orthologous genes, or panorthologs, that are shared by all 60 strains in the *E. coli* collection using the pipeline described in Cooper *et al.* (2010). To guard against recent gene duplication or horizontal transfer events, we confirmed that none of these panorthologs had better local BLAST hits in any given genome. We refer to these panorthologs as core genes, and to other genes that are present in only some of the *E. coli* collection as flexible genes. We realize that several strains in this collection are, to varying degrees, redundant; nonetheless, our findings are robust. We reran our analyses on a non-redundant subset of 15 genomes in the *E. coli* collection (NC_000913, NC_002695, NC_011415, NC_011601, NC_011745, NC_011750, NC_011751, NC_012967, NC_013353, NC_013654, NC_017634, NC_017641, NC_017644, NC_017663, NC_018658). Sequence diversity estimates from the complete *E. coli* collection are lower than estimates from the non-redundant subset of 15 genomes, as expected. The core genome of the full *E. coli* collection is also smaller at 1968 genes, in contrast to 2656 for the non-redundant subset of 15 genomes, which justifies the use of the complete collection in identifying a more tightly constrained set of core genes.

The NCBI Refseq accession for the ancestor for the LTEE, *E. coli* B strain REL606, is NC_012967. The accession for the *S. enterica* strain used as an outgroup is NC_003197. We downloaded *E. coli* and *S. enterica* orthology information from the OMA orthology database (Altenhoff *et al.* 2015), examining only the one-to-one matches. For internal consistency, we also used the panortholog pipeline to generate one-to-one panorthologs between *E. coli* B strain REL606 and *S. enterica*. We analyzed the 2853 panortholog pairs that the pipeline and the OMA database called identically.

### Analysis of the Keio Collection

We downloaded data on essentiality and growth yield in rich and minimal media for the Keio collection of single-gene knockouts in *E. coli* K-12 from the supplementary tables in the paper describing that collection (Baba *et al.* 2006). We classified the knocked-out genes as panorthologs (i.e., core) or not (i.e., flexible), and we compared differences in essentiality and growth yield between the two sets of genes.

### Nonsynonymous and Synonymous Mutations in the LTEE at 50,000 Generations

We identified all mutations in protein-coding genes in the whole-genome sequences of single clones isolated from each of the 12 LTEE populations at 50,000 generations. These 12 genomes are among the 264 genomes from various generations described by Tenaillon *et al.* (2016). Six of the 12 populations descend from REL606, and six from REL607 (Lenski *et al.* 1991). These ancestral strains differ by point mutations in the *araA* and *recD* genes (Tenaillon *et al.* 2016), and those mutations were thus excluded from our analysis. These 12 independently evolved genomes were used specifically in the initial categorical analyses reported in the section on “Core Genes Evolve Faster than Flexible Genes in the LTEE.”

### *G* Scores and Positive Selection on Genes in the LTEE

We use the *G*-score statistics reported in Supplementary Table 2 of Tenaillon *et al.* (2016) as a measure of positive selection at the gene level in the LTEE. The *G* score for each gene reflects, in a likelihood framework, the number of independent nonsynonymous mutations in nonmutator lineages relative to the number expected given the length of that gene’s coding sequence (relative to all coding sequences) and the total number of such mutations. In this analysis, the nonmutator lineages included the six LTEE populations that never evolved point-mutation hypermutability as well as lineages in the other populations before they became mutators. This analysis used the whole-genome sequences from all 264 clones isolated at 11 time points through 50,000 generations of the LTEE; only independent mutations were counted, but they were not necessarily present in the 50,000-generation samples.

### Sequence Diversity and Divergence

We adapted Nei’s nucleotide diversity metric (Nei and Li 1979) for use with amino-acid sequences to reflect nonsynonymous differences. Specifically, we calculated the mean number of differences per site between all 1770 (i.e., 60 x 59 / 2) pairs of sequences in the protein alignments from the 60 genomes in the *E. coli* collection. We counted each site in an indel between two sequences separately, so an indel that affected 10 amino-acid residues would count as 10 differences, even though it was probably caused by a single mutational event. In the site-specific analysis, we calculated this diversity metric separately for the sites that evolved in the LTEE and those that did not, and we compared the values to see if the former also tended to vary in nature. For the sequence divergence between *E. coli* and *S. enterica*, we used the ancestral strain of the LTEE, REL606, as the representative *E. coli* genome in order to maximize the number of orthologous genes available in our analysis. The divergence for each gene was calculated as the proportion of amino-acid residues that differ between the two aligned proteins, where an amino-acid difference implies at least one nonsynonymous change in the corresponding codon since the most recent common ancestor of the two alleles.

### Mapping Mutations to X-ray Crystal Structures

The full-length amino-acid sequence of 15 proteins (*topA, spoT, nadR, atoC, infC, rpsD, hflB, yijC, pykF, rpoB, mrdA, rne, ftsI, rplF, infB)* were aligned using Jackhmmer (HMMER version 3.1b2) against the PDB sequence database at www.rcsb.org (Berman *et al.* 2000, downloaded August 29, 2016). Mutated residues were visualized on the structure that had the best hit to the amino-acid sequence. In those cases where the best available structure was not from *E. coli,* we extracted the correct residue numbering from the Jackhmmer alignment.

### Evolutionary Tree Construction

We aligned all 1759 panorthologs shared between the *E. coli* collection and *S. enterica* using MAFFT (Katoh and Standley 2013), and we counted the amino-acid differences between all pairs of genomes using Biopython (Cock *et al.* 2009). We then used this distance matrix to construct a neighbor-joining tree (Saitou and Nei 1987). We affixed the 12 nodes for the 50,000-generation LTEE clones to the node of their ancestor, REL606, in the tree; their branch lengths are the number of nonsynonymous mutations in the 1759 panorthologs in each LTEE clone. We visualized the tree using the ETE Toolkit for phylogenomic data (Huerta-Cepas *et al.* 2016).

### Computational and Statistical Analyses

All data tables and analysis scripts will be deposited in the Dryad Digital Repository upon acceptance (doi: pending).

## Acknowledgments

We thank Alita Burmeister, Michael Wiser, and Kyle Card for comments on earlier versions of our manuscript; Jeff Barrick and Daniel Deatherage for making the LTEE genomics data accessible; and Thomas Hopf for assistance with computational analyses. This work was supported, in part, by a National Defense Science and Engineering Graduate Fellowship to RM, a grant from the National Science Foundation (DEB-1451740) to REL, the BEACON Center for the Study of Evolution in Action (NSF Cooperative Agreement DBI-0939454), and grants from NIGMS (R01GM106303) and the Raymond and Beverly Sackler Foundation to DSM.

---

Supplementary File 1. NCBI Refseq ID, strain name, and lifestyle (commensal or pathogen) for the collection of 60 strains with complete genome sequences used to identify the set of panorthologs that represent the core genome of *Escherichia coli*.

Supplementary File 2. UniProt and Pfam annotations for the 57 genes with two or more parallel nonsynonymous changes in nonmutator LTEE genomes, as reported in Tenaillon *et al.* (2016). The specific identity of the LTEE mutations found in these genes are provided for easy reference.

